# Pollinators in Mexico. Spatial patterns show exceptions to global trends

**DOI:** 10.1101/2025.10.30.685361

**Authors:** Angela Nava Bolaños, Luis Osorio Olvera, Jorge Soberón Mainero

## Abstract

Pollinators are declining globally, yet their biodiversity patterns in megadiverse regions remain unclear. We analyzed four major pollinator groups across one of the world’s most diverse regions –Mexico and adjacent areas. Using species distribution models, we generated presence– absence matrices (1,135 species in 165,078 grid cells) and applied range–diversity plots with a new randomization approach, to explore richness and endemism. Results reveal sharp contrasts: hummingbirds, bats, hawkmoths, and some bee families show the expected tropical richness peak, whereas other bees peak in arid lands of northern Mexico and the southwestern US. Endemism also diverges, with vertebrates concentrated in the tropics and bees in deserts. These findings challenge assumptions of uniform pollinator diversity patterns and demonstrate that conservation cannot rely on one-size-fits-all solutions. Our framework offers spatial tools to identify biodiversity structure and inform conservation and policy in the face of land-use change, agrochemicals, invasive species, and climate pressures.

## Introduction

Pollinators are essential for ecosystems and agriculture, contributing to over 70% of global crops (Klein *et al*. 2007). Their annual economic value has been estimated at $170 US billion (Gallai and Vaissière 2009). Yet pollinators are declining worldwide and are a global concern (Potts *et al*. 2016). To preserve pollinators, documenting their biodiversity patterns is both indispensable and challenging, given the information shortfalls prevalent (Hortal *et al*. 2015). Mexico is a megadiverse country and the second most specious country for bees in the world (Orr *et al*. 2021). However, spatial patterns of its pollinators remain poorly known (Nava-Bolaños *et al*. 2023). It is not even known whether pollinators in Mexico follow general trends, like the increased number of species towards the tropics, nor where is endemism concentrated. Here we document large-scale spatial patterns of biodiversity for Mexico’s major pollinators (bees, hawk moths, hummingbirds, and pollinating bats). We use relatively new tools to calculate biodiversity indices, using Presence-Absence Matrices (PAM) (Arita *et al*. 2012) and statistics derived from them (range-diversity plots). These innovative methods can be applied to other taxa and regions, offering insights needed for pollinator conservation decisions, urgent in the current context of loss of diversity of these organisms and their ecological service on which our food depends.

## Methods

### Input data

We compiled a database for Mexico’s main pollinator groups (bees, hawkmoths, hummingbirds, and glossophagine bats). Species names (2,404) were obtained from “*El Capital Natural de México”* (Sarukhan 2008). Occurrence records (~1.7 million) were downloaded from GBIF (https://doi.org/10.15468/dd.6ajy7v; accessed March 2020) using the R package spocc, and cleaned with ntbox (Osorio-Olvera *et al*. 2020) to remove duplicates, taxonomic and geographic inconsistencies, and records prior to 1950, following the workflow described in (Nava-Bolaños *et al*. 2023). The cleaned dataset comprised 216,058 unique records for 1,709 species. Environmental predictors included 15 bioclimatic variables from WorldClim (0.0416° resolution).

### Ecological Niche Models

Minimum□volume ellipsoids (MVEs) were fitted to occurrences of each species (Qiao *et al*. 2016), shape of MVEs is consistent with fundamental niches (Drake 2015), and fitting requires significantly fewer arbitrary decisions than other methods. Although MVE is a presence-only algorithm, ntbox uses environmental background data for estimating: 1) background prevalence of species; 2) area under the curve (AUC) of receiver operating characteristic curve (ROC); 3) partial ROC; and 4) binomial test. We extracted environmental information for records for each species and removed environmental duplicates. Datasets were split randomly into training and testing data in a proportion of 70:30. We estimated correlations among environmental variables (threshold of *r* > 0.7). MVE models were calibrated with ellipsoid_selection function. Ellipsoid models were built in 3, and 4 dimensions, using all possible combinations of the 15 least correlated variables. Models were run in parallel. The selected subset of the best models for each species are detailed in table “Metrics of Performance of Ellipsoid Fitting, per species” at the FigShare DOI https://doi.org/10.6084/m9.figshare.c.6651671.v1. This table contains metrics of performance of fit for 1,135 pollinator species. It includes columns for Order, Family, Species names, Bioclim variables used, Number of variables used (i.e., 3 means a 3-dimensional ellipsoid), omission rate predicting training points, omission rate predicting testing points, background prevalence, and the AUC ratios.

### PAM and biodiversity analysis

Models were clipped to “Megamexico” (Rzedowski 1991) (Mexican territory and border ecoregions). This prevents biasing the description of richness, endemism, and biogeographic processes by political borders. We clipped model projections with functions crop and mask from R raster package. Then, we overlap these Megamexico maps to build the pollinator PAM.

PAM are mathematical representations of lists of species in places. The use of PAM allows us to visualize biodiversity patterns in a detailed way, based on indices that can be calculated from them (Soberon and Cavner 2015). For example, in a sites by species PAM, the sum of the values in rows produce the number of species present in a given site, and the sum of the values in columns count in how many sites a species is present. Therefore, the column sums approximates the size of the distribution area in units of cell size and the row sums gives us an estimate of richness. A diversity-range plot (Soberón and Ceballos 2011) use these simultaneously. Diversity-range plot shows jointly two indices that describe the species composition in each cell of a map: (1) richness in the number of species, and (2) dispersion field, which refers to the amount of species overlapped in a cell with respect to the other cells (Soberón and Ceballos 2011). Diversity-range compacts data in an easy to interpret way. Furthermore, indices in the diversity-range plot have a simple geographic interpretation (Soberón and Ceballos 2011). Details of these calculations and equations can be found in Arita *et al*. 2008; Soberón and Ceballos 2011). These analyses are programmed and optimized for use in large geographic areas in the ‘bamm’ package with the function diversity_range_analysis. The returned object of this function includes the following rasters: alpha diversity, dispersion field, and diversity range, and an interactive diversity range plot. These analyses were performed for all pollinator species, as well for the 4 orders included, and finally for each pollinator family of Apoidea.

To estimate uncertainty of the metrics derived from a PAM, it is necessary to randomize it fixing the marginal values. This is a numerically challenging problem (Soberón *et al*. 2021), and we use an algorithm proposed by Strona et al. 2014. The method is implemented in the R ‘bamm’ package and runs very efficiently, allowing marginal-fixed randomization of matrices with tens of millions of cells. With fixed-marginals, the dispersion field value is randomly generated. We distinguished the two tails, and define as random values between the lowest and highest decils.

A schematic workflow summarizing full methods is provided in Supplementary Material (SM) Fig. 1, and summary of occurrence datasets, PAM, and model performance metrics are available at FigShare (https://doi.org/10.6084/m9.figshare.c.6651671.v1).

**Fig. 1:**
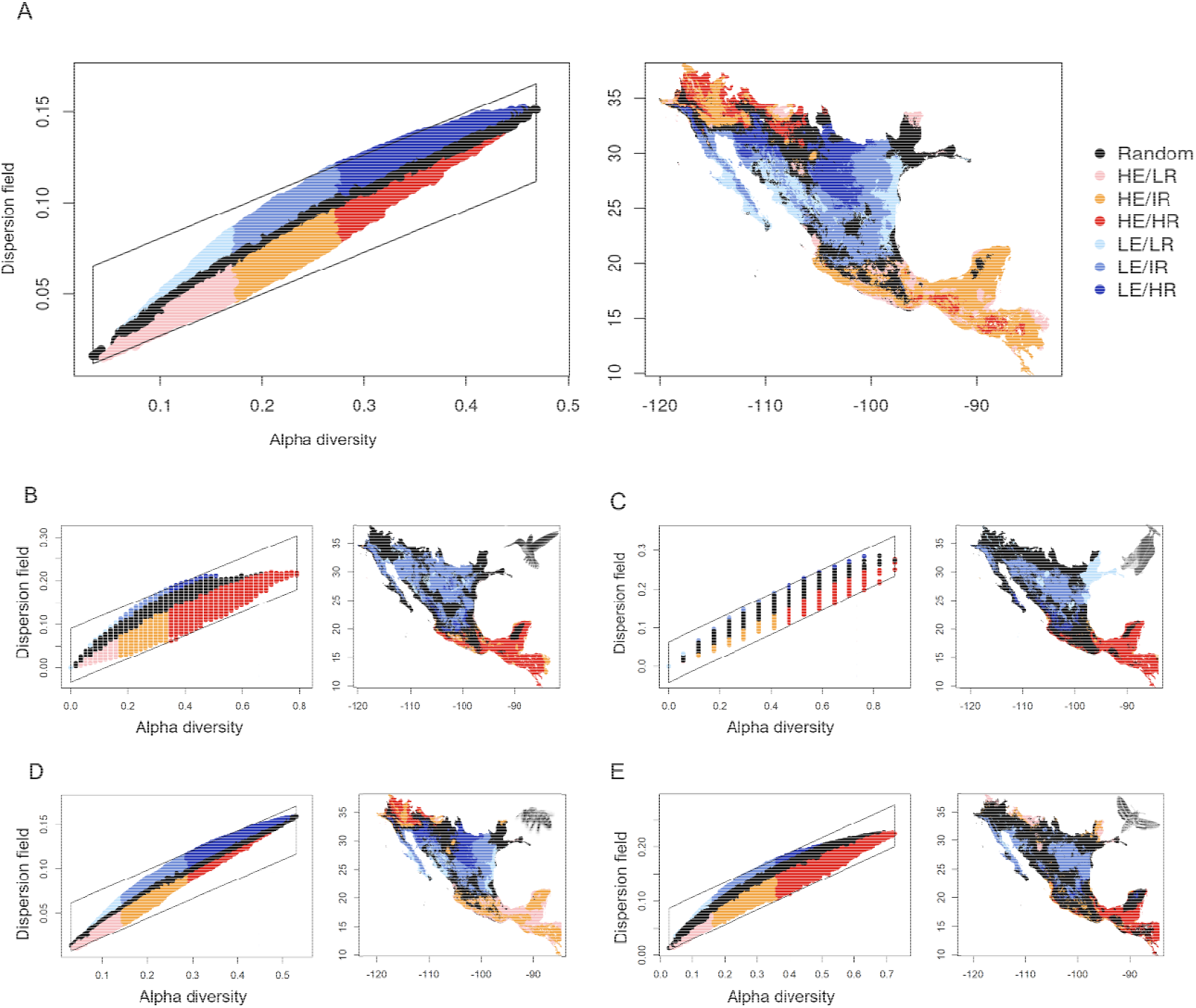
Range diversity patterns for pollinators. In A are presented the patterns for all species of pollinators, in B for hummingbirds, in C for bat pollinators, in D for bees, and in E for moths, in the left are the range diversity patterns presented in plots and in the right their geographical representations. Regions with high endemism and high richness (HE/HR) are represented in red, regions with high endemism and intermediate richness (HE/IR) are presented in orange, regions with high endemism and low richness (HE/LR) are presented in salmon, regions with low endemism and high richness (LE/HR) are presented with a stronger blue, regions with low endemism and intermediate richness (LE/IR) are presented with a medium blue, regions with low endemism and low richness (LE/LR) are presented with a light blue, and black are those regions which are not different to a random patterns.

## Results

### General patterns

Sampling of pollinators was highly uneven (SM Fig. 2), with taxonomic and spatial biases. We used distribution modelling to compensate for the bias (Lira-Noriega *et al*. 2007), and built a PAM with the results of the modelling (Vollering *et al*. 2016; Borregaard *et al*. 2020). We focused on two indices derived from the 165,078 (grid cells) X 1,135 (species) PAM: local (to cells) number of species (alpha diversity) and the “dispersion field” (Borregaard *et al*. 2020), which is equivalent to the mean similarity of a cell to all others (Soberon and Cavner 2015), and thus inversely related to endemism. Mapping the values of these indices reveal important differences among taxa [Figs 1,2], with a non-intuitive pattern of high richness and high endemism in the arid lands of North America, for most families of bees, and an opposite pattern of tropical high richness and high endemism for hawkmoths and vertebrates.

**Fig. 2:**
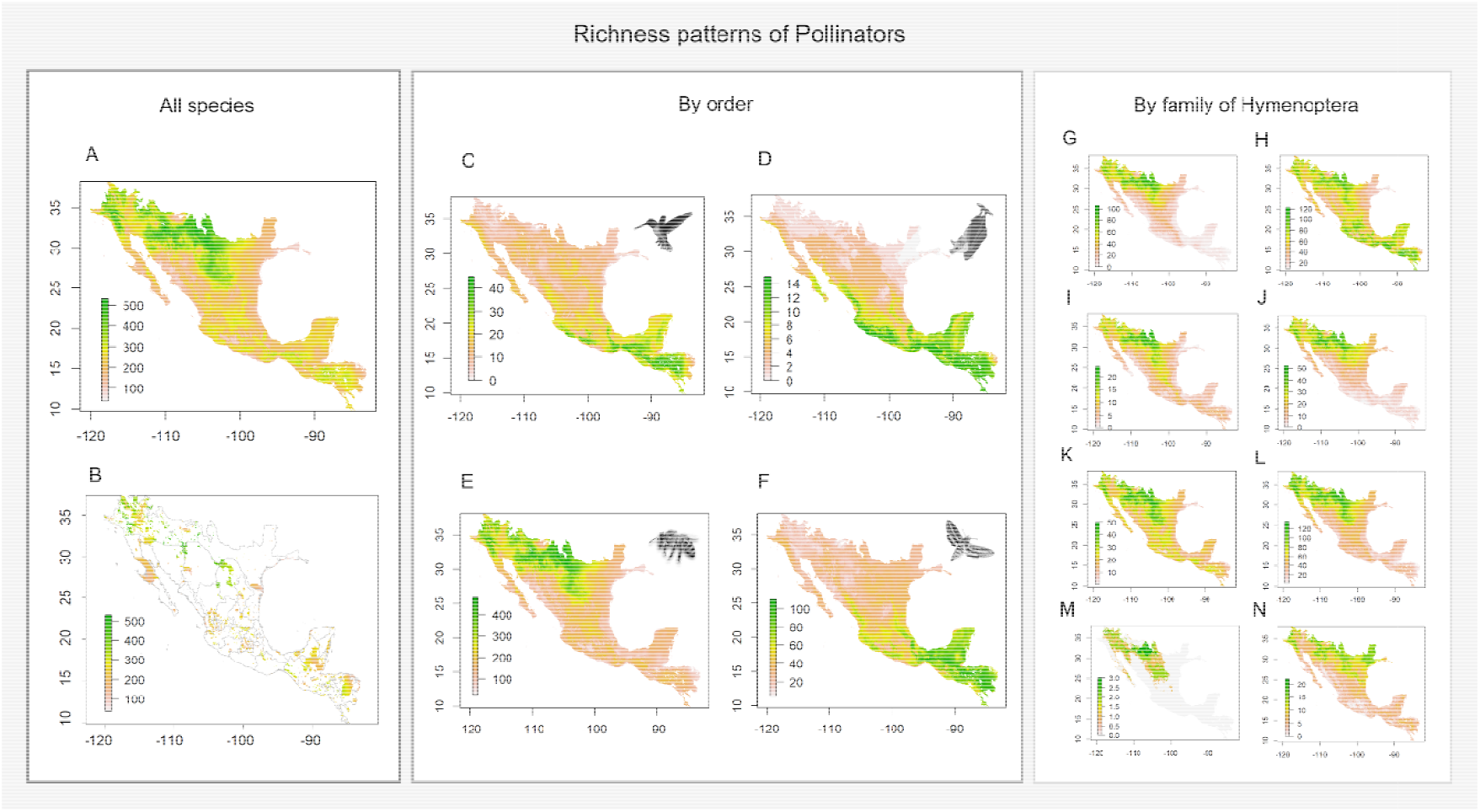
Alpha diversity patterns for pollinators. In A are presented richness patterns for all pollinators species, in B richness patterns presented in all Protected Nature Areas for the region of Megamexico. From C to F richness patterns are presented by order of pollinators: In C for hummingbirds, in D for bat pollinators, in E for bees and in F for moths. From G to N richness patterns are presented by family of Apoidea: In G for Adrenidae, in H for Apidae, in I for Colletidae, in J for Crabronidae, in K for Halictidae, in L for Megachilidae, in M for Melittidae, and in N for Sphecidae.

How different are patterns from what would be obtained from different classes of null and neutral models? One way to answer this question is by randomizing the PAM subject to different constraints. For null models, randomizing large PAM when both the marginals are kept constant is not a simple problem. We present software to do this, based on an algorithm described by Strona et. al. (Strona *et al*. 2014).

Randomized range-diversity plots (Soberón *et al*. 2021) highlight non-random geographic patterns [Fig. 1]. Geographically, regions of high-richness and high endemism regions tend to be separated from those of low richness, low endemism, by zones with patterns indistinguishable from random. Since the richness patterns are not randomized, the randomness pattern displayed in Fig. 1 is due to the “dispersion field”, which is an index of similarity among cells in the grid.

The geographic pattern displayed in Fig. 1 shows that most of the taxa (Apidae) display bimodal endemicity: in northwest and south Megamexico. However, endotherms (including hawkmoths, see discussion) display patterns that differ from the bees. In the interactive graphic, the correspondence between different regions in the plot and in the map can be interactively explored (SM Fig. 3), looking for statistically significant combinations of richness and endemism.

**Fig. 3:**
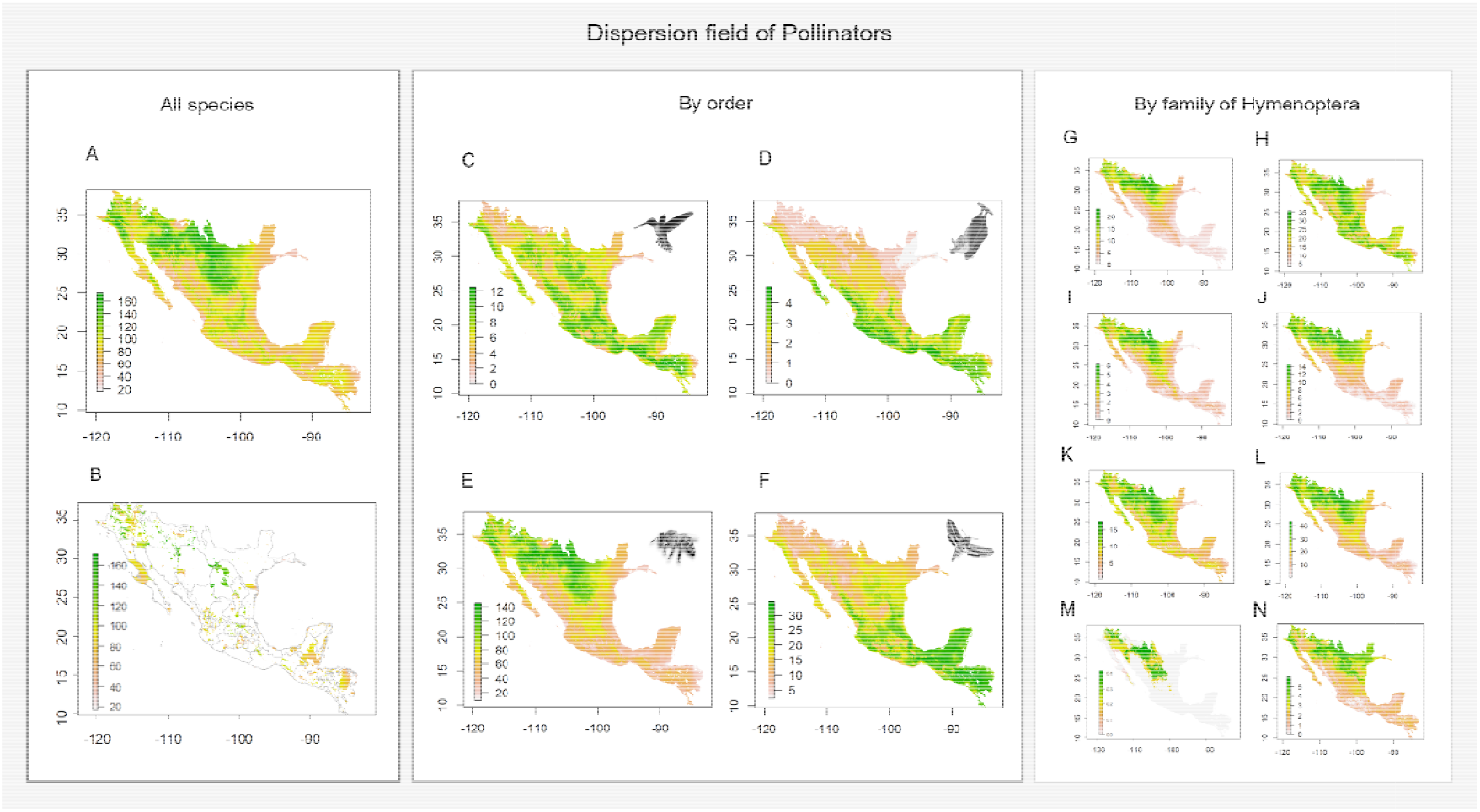
Dispersion field patterns for pollinators. In A are presented dispersion field patterns for all pollinators species, in B dispersion field patterns presented in all Protected Nature Areas for the region of Megamexico. From C to F dispersion field patterns are presented by order of pollinators: In C for hummingbirds, in D for bat pollinators, in E for bees, and in F for moths. From G to N dispersion field patterns are presented by family of Apoidea: In G for Adrenidae, in H for Apidae, in I for Colletidae, in J for Crabronidae, in K for Halictidae, in L for Megachilidae, in M for Melittidae, and in N for Sphecidae.

### Patterns by group

The highest richness in Megamexico is found in bees (Apoidea), in the arid lands and deserts of NW Mexico and SW United States. A similar pattern has been previously reported for the bees of North America (Orr *et al*. 2021). The patterns vary by family (Fig. 1–3), and the spike of very high proportional richness associated with relatively low mean ranges is not general. In SM Table 1 we show the percentage of points in the grid that appear in each class of randomization, for species in the different orders. What is the meaning of these results? Randomization affects the compositional similarity (also called the dispersion field, which is inversely related to endemism), but not the richness. For all groups considered together, in 23.18% of the focal region, compositional similarity is indistinguishable from that obtained from a random permutation of the PAM, with a 38.89% of the territory being less-endemic than random and the rest more endemic than random. However, for the bees (H column SM Table 1), almost 37.5% of Megamexico is of significantly high endemism (HE/LR, HE/IR, HE/HR). For bees, only 4.5% of Megamexico is low in both endemism and richness. Regions with dispersion fields higher than random (i.e., regions of low endemism) tend to concentrate in the central part of the region, and those with endemism values higher than random, concentrate in the south and north. For the other groups, the regions indistinguishable from random tend to be in the central and northern parts, and the high endemism and high richness is a tropical pattern (most marked in the Hawkmoths).

## Discussion and Conclusion

### General patterns

It has been previously reported that, at a global scale, bees are an exception to the general pattern (Gaston 1996; Orr *et al*. 2021) of more species towards the equator (Bystriakova *et al*. 2018), presenting a higher number of species at medium latitudes. We confirm the temperate pattern for most families of bees in our region, although the Apidae (Fig. 2 H), that contains very tropical subfamilies (Euglossinae, Trigoninae and Meliponinae), is an exception. The pattern of more species in tropical areas hold for the bats, hummingbirds and hawkmoths. The highly correlated numbers of species among hummingbirds, bats and hawkmoths suggest a common causal factor, perhaps related with physiological responses to climate. Vertebrates are homeotherms, and Hawkmoths have behavioural ways for managing low temperatures. However, large bee families depart markedly from this pattern. This has the immediate implication that it is not very useful to generalize about the distributions of “pollinators” in general: not only the plants they pollinate, and their efficiencies as pollinators are different (Ne’Eman *et al*. 2010), but also the biogeographic patterns pollinators display.

Our method allows us to study patterns of endemism, because “mean dispersion field”, by definition, is the average of the ranges of distribution of the species living in a place. It can be seen as a measure of the size of the species pool and used to display geographic organization of sets of species (Borregaard *et al*. 2020). A large value of the dispersion field of a cell in the grid means cells occupied by geographically widespread species, (a non-endemic fauna), and vice versa. Our results show that, in our study region, zones of high number of species and restricted ranges occur mostly in the tropical part for the bats, hawkmoths and hummingbirds, with the bees concentrating both high richness and narrow distributions in north-western Mexico and the south-western United States. With exception of bees, the peninsula of Yucatan tends to have combinations of richness and endemism indistinguishable from a random permutation of the PAM. Again, with exception of bees, most of the Nearctic part of Mexico has low richness and low endemism.

Our randomization procedure fixes the richness and incidence marginals, a procedure consistent with the hypothesis that both niche and movement factors affect diversity (Gibert and Escarguel 2019). Whether climatic tolerances or dispersal capacities determine the pattern is a question for future research, but our contrasting results for different groups suggest that the relative importance of dispersal vs. niche preferences will be also different.

### Relevance of the methods

Previous similar work (Bystriakova *et al*. 2018; Orr *et al*. 2021) have used raw occurrence data to describe patterns. This approach is seriously affected by sampling incompleteness (Lira-Noriega *et al*. 2007). Our method uses niche-based species distribution models to compensate for the sampling biases, as in (Vollering *et al*. 2016). The PAM thus created are “potential distribution” PAM, which approximate a true PAM, based on actual distributions.

The use of range-diversity plots allows partitioning geography in regions. We split the plot in quartiles (see Fig. 1), corresponding to high, low and medium values on richness and similarity (dispersion field), and a region indistinguishable from random. Partition in the range-diversity plot has a geographical correspondence, clustering regions by richness and similarity (Fig. 1). This means that in some sense our partitioning method is an alternative to clustering algorithms like k-means, or k-medoids (Vollering *et al*. 2016). Our method is based on biological rather than geometric considerations, and therefore may provide deeper biological insight.

### Future directions

We constructed a first approximation to PAM for several groups of pollinator species in a region of North America (as defined above). A PAM is a very informative proxy for “biodiversity”, as we show, demonstrating different richness and endemism patterns for different groups. However, model-based PAM are dependent of two major assumptions: widespread dispersal capacities, and unimportance of biological interactions. Evaluating how important these assumptions are is the first task for the future. Our preliminary results suggest that ignoring these factors create mostly commission errors (overprediction).

The methods presented in this paper can be replicated for any other area in the world, as well as, for other taxa of pollinators. These analyses are necessary to make pollinator conservation decisions, which are necessary in the current context of loss of diversity and abundance of pollinators. Some of the future directions for which the results presented in this work can be used as input are for example: 1) evaluating the impact of climate change on richness and endemism patterns of pollinators, 2) evaluating the degree of protection offered by Protected Natural Areas for pollinating species under these dynamic scenarios, 3) study the different threats faced by species on a spatial and geographical scale, such as the industrial use of agrochemicals, invasive species, change in land use, human footprint, 4) there is a urgent necessity to promote research on pollinators in regions that remain unexplored and for species poorly studied, and 5) design of public policies that guarantee the conservation of these important species, which will largely guarantee food security and the human right to a healthy environment.

Although more work is needed, the bimodal latitudinal gradient is strongly supported, consistent with other studies which reported this richness pattern for bees in the world. Since there is no evidence suggesting a bounty of undescribed species that may change these principal patterns, our work supports a bimodal latitudinal pattern for bees.

## Acknowledgments and funding sources

To Secretaría de Educación, Ciencia, Tecnología e Innovación de la Ciudad de México (SECTEI), and Secretaría de Ciencia, Humanidades, Tecnología e Innovación (SECIHTI) for the support to ANB. To Exequiel Ezcurra, of the University of California at Riverside, for insightful comments on an early version of the work. LO-O acknowledges partial financial support from the UNAM Research and Technological Innovation Support Program (PAPIITIA202824) and the SECIHTI Frontier Science Project(CF-2023-I-115). LO-O also thank Nancy Gálvez-Reyes, Graciela García-Guzmán,and Miguel Baltazar Gálvez for their technical support in projects IA-202824 and CF-2023-I-115.

## Supplementary Materials

**Supplementary Material Figure 1.**
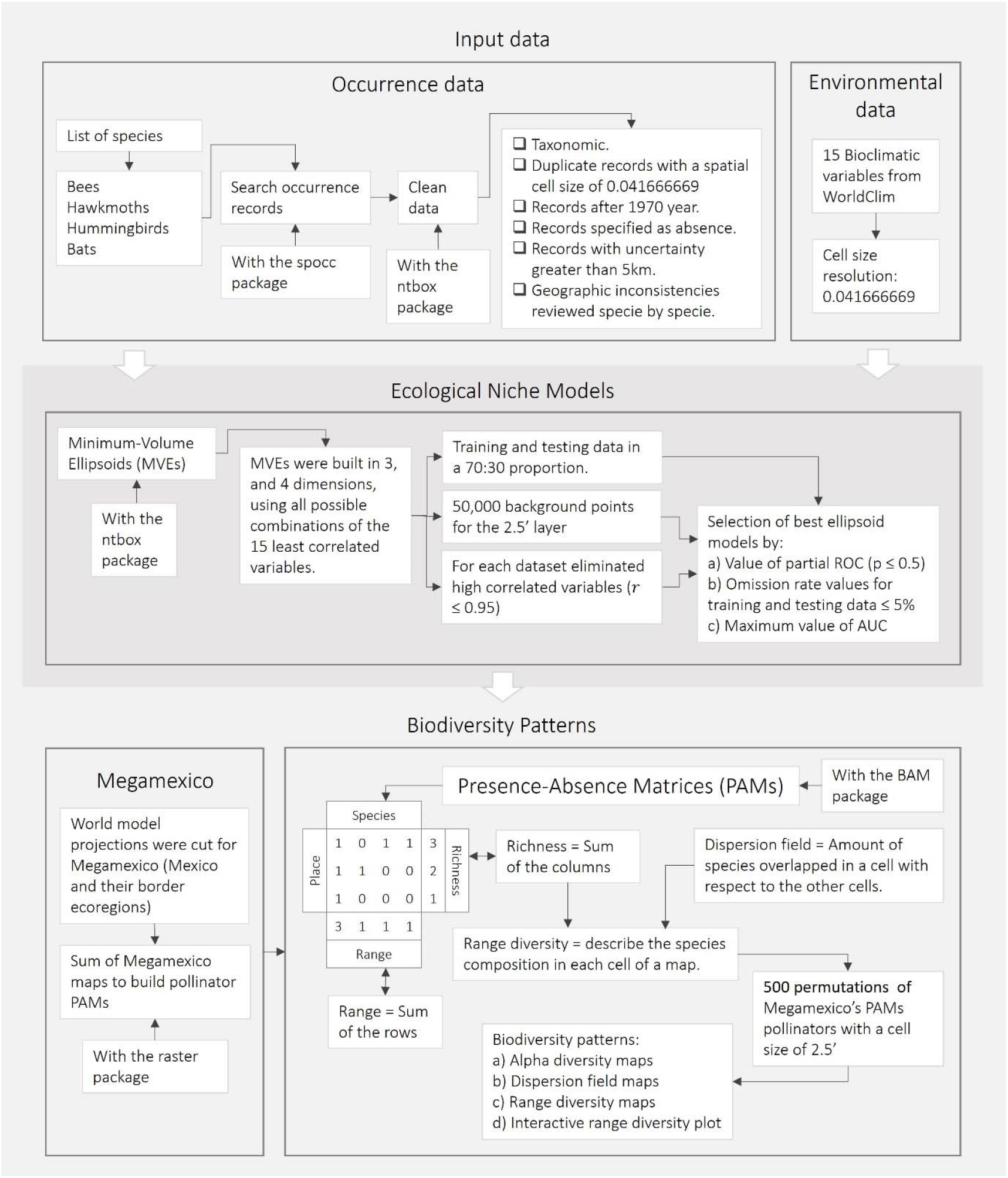
Workflow of methods. Here is a schematic representation of the steps to obtain patterns for pollinators in Megamexico, performing ecological niche models, analysis from PAM.

**Supplementary Material Figure 2.**
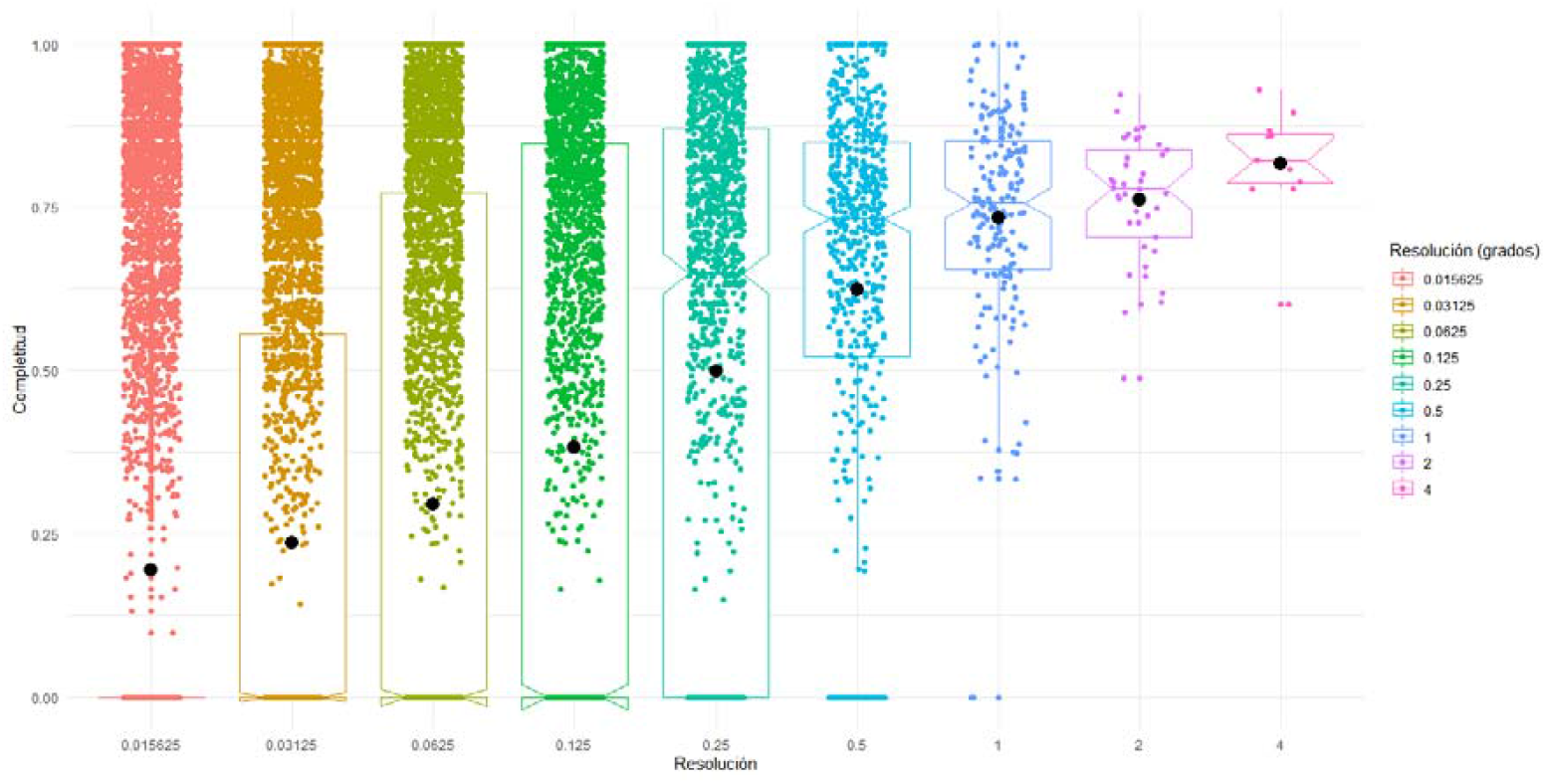
Index of completeness of inventories at different resolutions, in geographic degrees: It is well known that observation records are highly biased. To assess the bias in our dataset, we performed a completeness analysis of the database, for different resolutions of the data. Following (Soberón *et al*. 2007) we used a completeness of inventory index that runs from 1 (when increasing sampling effort fails to increase a list) to low numbers (when increases in effort linearly increase the size of the list). The results indicate that degree of completeness of the GBIF database we are using are reduced when resolution increases. Since we are aiming at an analysis at resolution of grid cells between 2.5 and 10 minutes of arc (~5km to 20km of side) it is clear that at these resolutions completeness of inventories is far from satisfactory. Usage of niche-based SDMs has been shown to compensate for this problem, assuming that the proper procedures are taken to compensate for over-sampling in certain localities.

Three large tables (Occurrence data, PAM, and metrics of performance of fit to ellipsoids) are deposited here https://doi.org/10.6084/m9.figshare.c.6651671.v1

**Supplementary Material Table 1.**
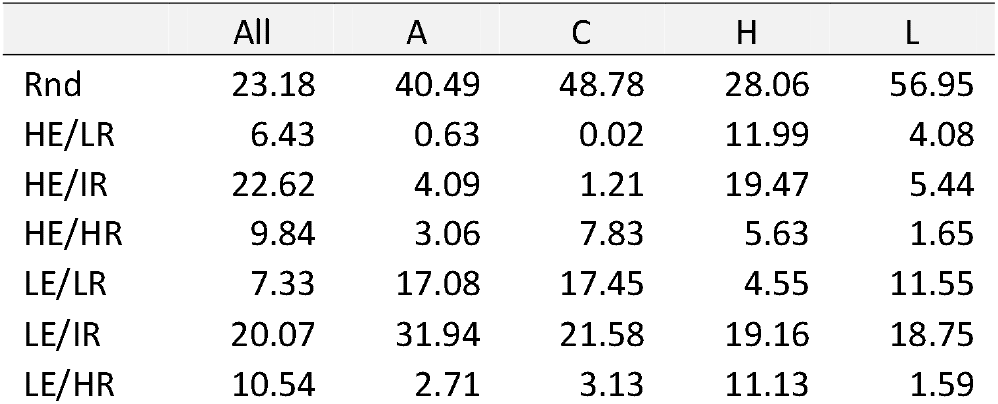
Table 1. Percentage of cells in the grid for which values predicted of dispersion field are indistinguishable from random (first row), significantly smaller than random (lower 10% quantile, HE rows), and significantly larger than random (upper 90% quantile, last three rows, LE). Richness in the lowest quartile of the distribution of is labelled LR, in the highest quartile is labelled HR, and in between IR. A=Hummingbirds, C= Bats, H=bees and L=Hawkmoths, in All species in all groups are pooled.

